# Innolysins: A novel approach to engineer endolysins to kill Gram-negative bacteria

**DOI:** 10.1101/408948

**Authors:** Athina Zampara, Martine C. Holst Sørensen, Dennis Grimon, Fabio Antenucci, Yves Briers, Lone Brøndsted

## Abstract

Bacteriophage-encoded endolysins degrading the essential peptidoglycan of bacteria are promising alternative antimicrobials to handle the global threat of antibiotic resistant bacteria. However, endolysins have limited use against Gram-negative bacteria, since their outer membrane prevents access to the peptidoglycan. Here we present Innolysins, a novel concept for engineering endolysins that allows the enzymes to pass through the outer membrane, hydrolyse the peptidoglycan and kill the target bacterium. Innolysins combine the enzymatic activity of endolysins with the binding capacity of phage receptor binding proteins (RBPs). As our proof of concept, we used phage T5 endolysin and receptor binding protein Pb5, which binds irreversibly to the phage receptor FhuA involved in ferrichrome transport in *Escherichia coli*. In total, we constructed twelve Innolysins fusing endolysin with Pb5 or the binding domain of Pb5 with or without flexible linkers in between. While the majority of the Innolysins maintained their muralytic activity, Innolysin#6 also showed bactericidal activity against *E. coli* reducing the number of bacteria by 1 log, thus overcoming the outer membrane barrier. Using an *E. coli fhuA* deletion mutant, we demonstrated that FhuA is required for bactericidal activity, supporting that the specific binding of Pb5 to its receptor on *E. coli* is needed for the endolysin to access the peptidoglycan. Accordingly, Innolysin#6 was able to kill other bacterial species that carry conserved FhuA homologs such as *Shigella sonnei* and *Pseudomonas aeruginosa*. In summary, the Innolysin approach expands recent protein engineering strategies allowing customization of endolysins by exploiting phage RBPs to specifically target Gram-negative bacteria.

**IMPORTANCE:** The extensive use of antibiotics has led to the emergence of antimicrobial resistant bacteria responsible for infections causing more than 50,000 deaths per year across Europe and the US. In response, the World Health Organization has stressed an urgent need to discover new antimicrobials to control in particular Gram-negative bacterial pathogens, due to their extensive multi-drug resistance. However, the outer membrane of Gram-negative bacteria limits the access of many antibacterial agents to their targets. Here, we developed a new approach, Innolysins that enable endolysins to overcome the outer membrane by exploiting the binding specificity of phage receptor binding proteins. As proof of concept, we constructed Innolysins against *E. coli* using the endolysin and the receptor binding protein of phage T5. Given the rich diversity of phage receptor binding proteins and their different binding specificities, our proof of concept paves the route for creating an arsenal of pathogen specific alternative antimicrobials.

## INTRODUCTION

The limited permeability of the outer membrane is a major obstacle for development of novel antimicrobials against Gram-negative pathogens preventing many compounds from reaching their intracellular targets (1). Bacteriophages (phages), viruses that infect bacteria, have naturally evolved mechanisms to overcome the outer membrane to infect their bacterial hosts (2, 3). In the first step of infection, phages bind to host cells and inject their genetic material across the outer and inner membrane of the bacterial cells into the cytoplasm. Also, during the final stage of the lytic infection cycle, phages produce proteins within the cell, which destroy the bacterial cell wall, leading to cell lysis. Thus, the molecular tools developed during phage evolution may be exploited to develop novel phage-based antimicrobials that are able to pass the outer membrane and to kill Gram-negative bacteria.

Phages recognize their host bacteria by binding to specific surface receptors that may be outer membrane proteins, lipopolysaccharides or components of bacterial capsules, pili and flagella (4). The adhesion specificity is mediated by receptor binding proteins (RBPs) that form fibers or spikes at the distal phage tail. A well-characterized RBP is Pb5, located at the tail tip of the phage T5, which specifically binds to the bacterial receptor FhuA during infection of the *E. coli* host (5, 6). FhuA is an outer membrane protein that actively transports siderophore-ferrichrome and allows *E. coli* to take up iron from the environment. The crystal structure of FhuA reveals two domains, a 22-stranded anti-parallel barrel that forms a hollow channel with eleven surface-exposed loops and a globular domain, known as the plug, which blocks the channel in its inactive state (7, 8). Previously it was shown that Pb5 of phage T5 binds irreversibly to the extracellular loop L4 and the plug of FhuA. In addition, it was proposed that FhuA acts as an anchor for phage T5 binding to *E. coli*, subsequently allowing the tail fiber composed of Pb2 to transverse the outer membrane and inject the DNA (9, 10).

During the last stage of phage infection, phages produce endolysins to lyse the bacterial cells and release progeny phages. Endolysins are hydrolytic enzymes that, after gaining access to the periplasm, degrade the peptidoglycan, leading to cell lysis (11). Endolysins may have different catalytic activities, depending on the bond that they target in the peptidoglycan, and are classified as glycosidases, amidases or endopeptidases. The endolysin encoded by phage T5 is a well characterized endopeptidase that hydrolyzes the bond between L-alanine and D-glutamic acid of the peptidoglycan in *E. coli* (12). While native endolysins have been successfully applied as antimicrobials against Gram-positive bacteria including *Staphylococcus aureus* and streptococci, the outer membrane of Gram-negative bacteria generally prevents access to the peptidoglycan layer (13). To enable killing of Gram-negative pathogens using endolysins, a number of engineering strategies have been proposed. Recently it was shown that the fusion of endolysins to polycationic or amphipathic peptides enables the endolysins to pass the outer membrane and kill multi-drug resistant *Pseudomonas aeruginosa* and *Acinetobacter baumannii* (14). Furthermore, recombinant T4 lysozyme carrying the binding domain of pesticin, targeting the outer-membrane protein FyuA, has been shown to kill *Yersinia* and pathogenic *E. coli* strains (15). Similarly, fusion of the *E. coli* phage endolysin Lysep3 with the translocation and receptor-binding domain of another bacteriocin colicin A, targeting the outer membrane protein BtuB could kill *E. coli* (16). Thus, these studies have demonstrated that endolysins can be engineered to pass the outer membrane and subsequently kill Gram-negative bacteria.

Here, we expand on the concept of delivering endolysins across the Gram-negative outer membrane by combining the specificity of phage RBPs with the antimicrobial activity of endolysins into RBP-endolysin hybrids (Innolysins). To provide a proof of concept, we utilize the endolysin and the RBP (Pb5) of phage T5. Pb5 was selected among many phage RBPs due to the specific and stable binding of Pb5 to its cognate protein receptor allowing us to target the fused phage T5 endolysin to the periplasm, where it can degrade the peptidoglycan. This engineering approach has the potential to utilize multiple distinct phage-derived RBPs to customize the bactericidal spectrum of Innolysins against diverse Gram-negative bacteria.

## RESULTS

### Strategy for construction of Innolysins

To construct Innolysins, we combined the RBP of phage T5 (Pb5) with the phage T5 endolysin (T5 Lys) for targeted delivery of the endolysin. The binding domain of Pb5 has previously been shown to be located in the N-terminus (488aa) of the protein (17). To determine whether the binding domain of the Pb5 is sufficient to enable the endolysin to pass through the outer membrane, we fused T5 endolysin with both the entire Pb5 and with the Pb5 binding domain (Pb5_1-488_). We anticipated that antimicrobial activity of an Innolysin requires both that Pb5 is able to bind to the outer membrane protein FhuA, and that the fused phage T5 endolysin is translocated across the outer membrane and remains active to degrade the peptidoglycan. Thus, to ensure that the joined domains remain functional after fusion, we either fused them directly or added linkers in between. We used flexible linkers composed of small non-polar amino acids, glycine and alanine, providing a certain degree of movement of the fused domains (18). To optimize the two-domain cooperation, we used linkers of two different sizes, L1 composed of six amino acids and L2 that consists of fourteen amino acids. In addition, to investigate the optimal orientation of the endolysin and RBP domains, we constructed the Innolysins in two directions with and without linkers (Fig. 1). As such, a total of twelve Innolysins were constructed.

**FIG 1:**
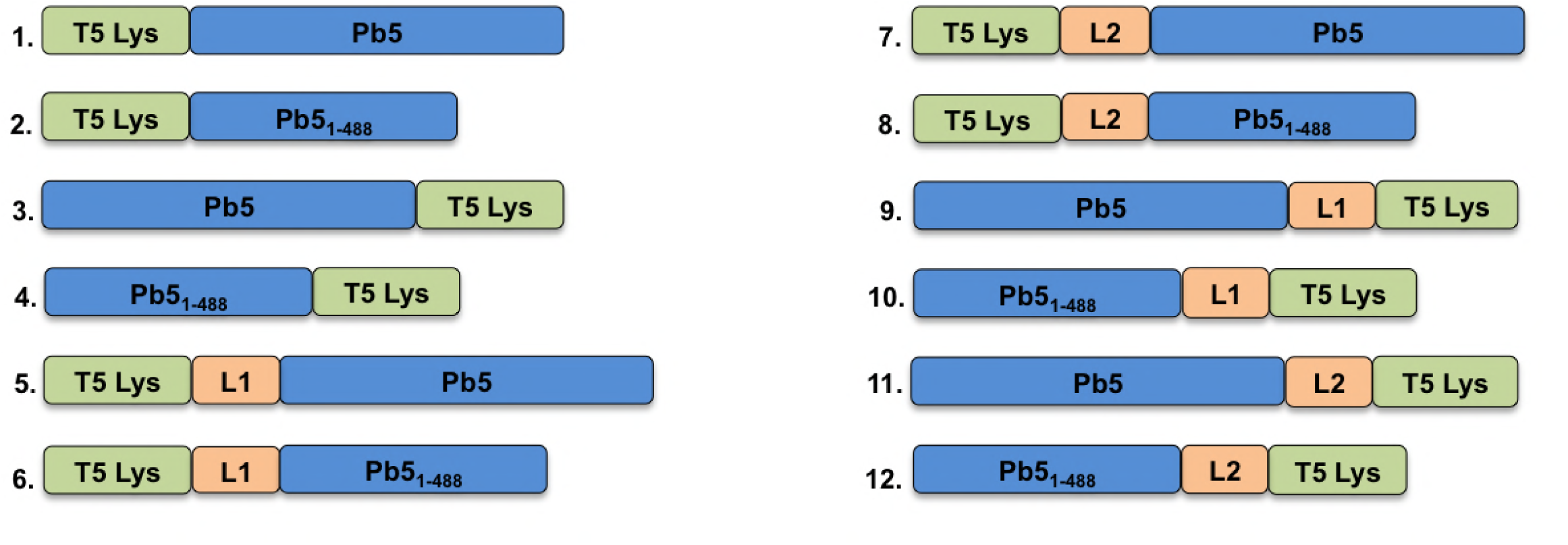
Visual representation of Innolysins. Phage T5 endolysin (T5 Lys) was fused with the whole phage T5 receptor binding protein, Pb5 or the binding domain of Pb5 (Pb5_1-488_). Fusion was conducted without linkers (1 - 4) or with linkers (L1 or L2) (5 - 12) with T5 Lys located in the N-terminus (1-2; 5 - 8) or C-terminus (3 - 4; 9 - 12) of the engineered proteins.

### Innolysins show muralytic activity

To demonstrate that the muralytic activity of the endolysin is maintained after the fusion with the RBP and a linker, the fused proteins were expressed in *E. coli* BL21 and supernatants of cell lysates were tested for muralytic activity. A standardized assay for analysis of the muralytic activity of endolysins acting against peptidoglycan of Gram-negative bacteria was used, based on outer membrane permeabilized *P. aeruginosa* cells, which share a common peptidoglycan chemotype (A1γ) with *E. coli* (19). The majority of Innolysins (nine out of twelve) were active with enzymatic activity ranging between 126-771 U/ml. Innolysin#9 (Pb5-L1-T5Lys) showed the highest activity per ml cleared lysate, reaching the muralytic activity of phage T5 endolysin alone (795 U/ml) (Fig. 2). Although expression was confirmed for all constructs, three of the Innolysins (#1, #3 and #12) and Pb5 alone did not show any significant activity compared to the negative control (muralytic activity of supernatant cell lysates carrying the empty vector, pVTSD2). Interestingly, both Innolysin#1 and #3 were composed of phage T5 endolysin and the whole Pb5 protein, fused in two opposite directions without a linker. Overall, the majority of hybrid endolysins showed muralytic activity after fusion with Pb5 either in the N-or C-terminus, while the presence of a flexible linker enhanced the activity of the engineered proteins.

**FIG 2:**
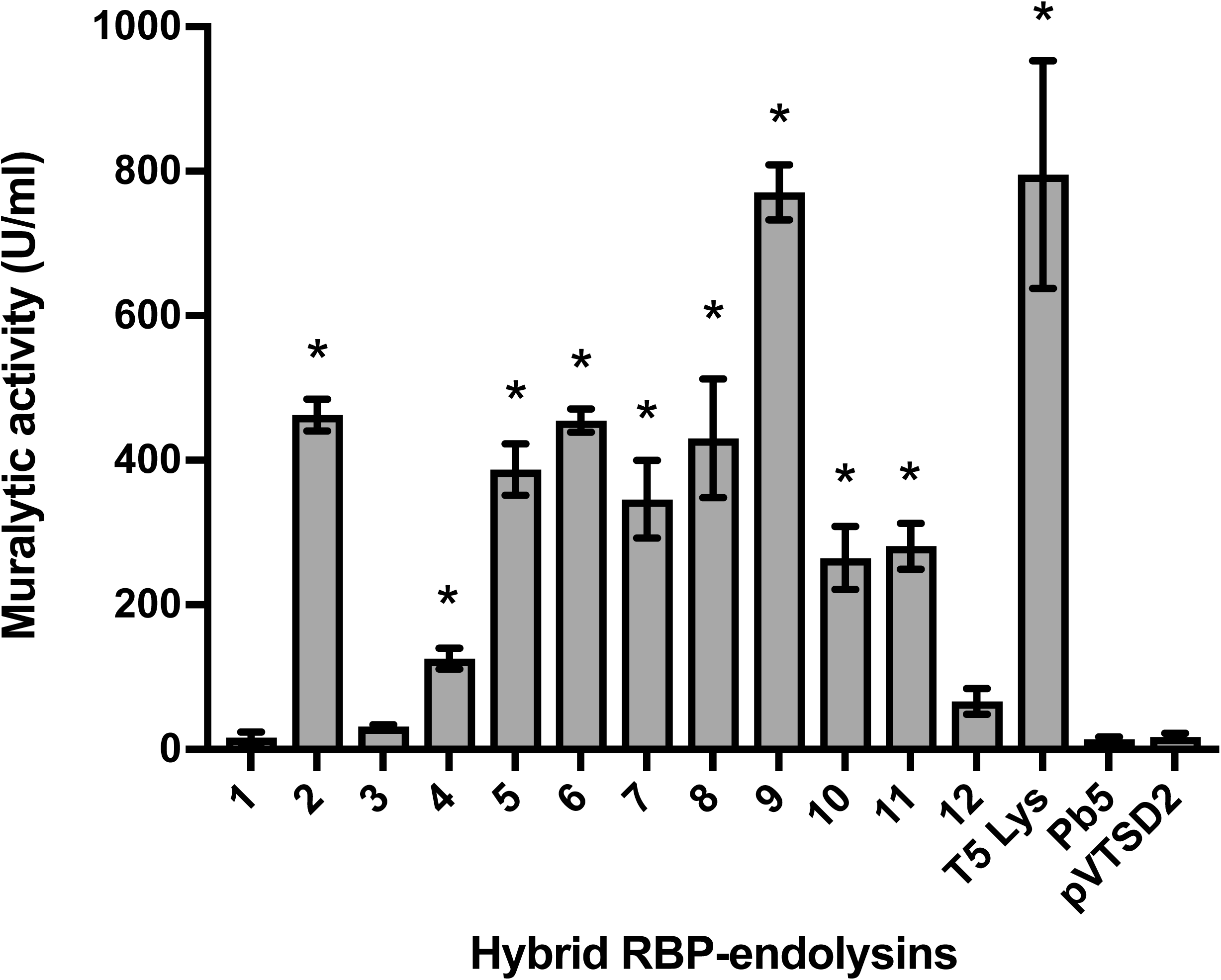
Muralytic activity of Innolysins. Soluble fractions of the cleared lysate of the engineered proteins (1-12, see Fig. 1) were screened for ability to degrade peptidoglycan of *P. aeruginosa* PAO1 and compared to the activity of soluble lysate of cells carrying the empty vector, pVTSD2. Phage T5 endolysin and receptor binding protein, Pb5 were used as a positive and a negative control, respectively. Average muralytic activities (U/ml) were estimated based on triplicates. * Significant muralytic activity at P < 0.05.

### Innolysin#6 inhibits *E. coli* growth

To assess whether binding of Pb5 allowed the fused endolysin to get access to the peptidoglycan, we screened the muralytically active Innolysins for their ability to inhibit growth of the phage T5 bacterial host *E. coli* ATCC11303. ATCC11303 was mixed with supernatants of cell lysates of strains expressing Innolysins, and bacterial growth was measured spectrophotometrically after 18h (Fig. 3). Growth inhibition was determined as the lack of growth of a start inoculum treated with an Innolysin compared to the negative control (growth of ATCC11303 cells treated with supernatant lysates of cells carrying the empty vector, pVTSD2). A significant inhibitory activity was noticed when *E. coli* ATCC11303 was treated with Innolysin#6 similar to the positive control, Art-175, an engineered endolysin that was previously shown to have both an inhibitory and bactericidal effect against *E. coli* (20, 21). The remaining eight Innolysins, Pb5 or endolysin alone did not significantly affect the *E. coli* growth compared to the negative control (Fig. 3). The screening resulted in one promising antimicrobial candidate out of the twelve Innolysins that could inhibit *E. coli* growth, indicating that domains have to be fused in a specific order and with a specific linker to acquire antibacterial activity after fusion. The active variant was composed of T5 endolysin in the N-terminus linked by a six amino-acid linker to the Pb5_1-488_ in the C-terminus. This result indicates that the binding domain of Pb5 alone can target the bacterial cells and allow phage T5 endolysin to overcome the outer membrane and inhibit *E. coli* growth.

**FIG 3:**
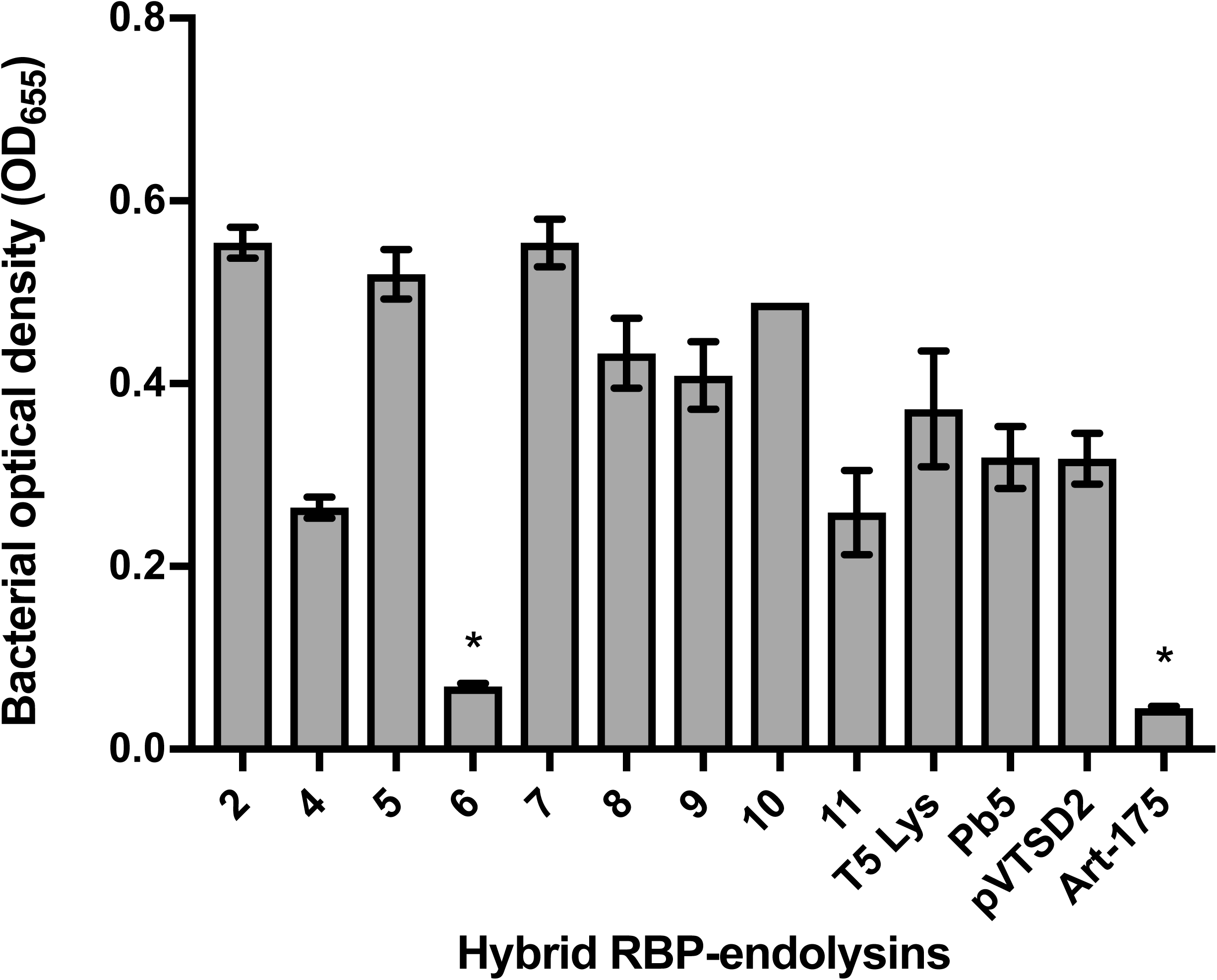
Growth of *E. coli* after treatment with Innolysins. Soluble induced fractions of muralytically active engineered proteins (see Fig. 2) were screened for the ability to inhibit growth of *E. coli* ATCC11303. Art-175 was provided by Lysando AG (20) and used as a positive control. Phage T5 endolysin and receptor binding protein, Pb5 were used as negative controls. Growth inhibition was detected as the absence of growth (OD_655nm_) of cells treated with an Innolysin in comparison to the negative control (growth of ATCC11303 cells treated with supernatant lysates of cells carrying the empty vector, pVTSD2) after incubation for 18 hours at 37°C. * Significant bacterial growth inhibition at P < 0.05.

### Killing efficiency of Innolysin#6 requires FhuA, but not energy

To determine whether binding of Pb5 to FhuA is required for the antimicrobial activity of Innolysin#6, killing efficiency of the purified hybrid protein was tested on a *fhuA* deletion mutant in *E. coli* BL21 and compared to the wild type *E. coli* BL21 and *E. coli* ATCC11303. Bacterial cells were incubated with the purified Innolysin at a final concentration of 0.2 mg/ml and the reduction in cell counts was compared to the non-treated cells (Fig. 4). *E. coli* BL21 cell counts were reduced by 1.22 ± 0.12 log after Innolysin treatment and a similar decrease was noticed in *E. coli* ATCC11303 cells (1.09 ± 0.15 log reduction), without prior optimization. No significant effect of the Innolysin was shown in *E. coli* BL21Δ*fhuA*, supporting our hypothesis that the Innolysin leads to peptidoglycan degradation when cells harbor the Pb5 receptor FhuA. While FhuA uptake of ferrichrome requires energy from the cytoplasmic proton motive force transduced to the outer membrane via the TonB protein, phage T5 interacts with FhuA independent of TonB (8, 22). To determine whether activity of Innolysin#6 requires such energy, bactericidal activity was tested against an *E. coli tonB* deletion mutant (ECOR4*ΔtonB*) and the wild type ECOR4 as a positive control (Fig 4). As expected, TonB was not required for bactericidal activity of the Innolysin (Fig. 4). Our combined data demonstrate that Innolysin#6 requires the presence of FhuA in order to be transported passively through the outer membrane and target the peptidoglycan, leading to killing of *E. coli*.

**FIG 4:**
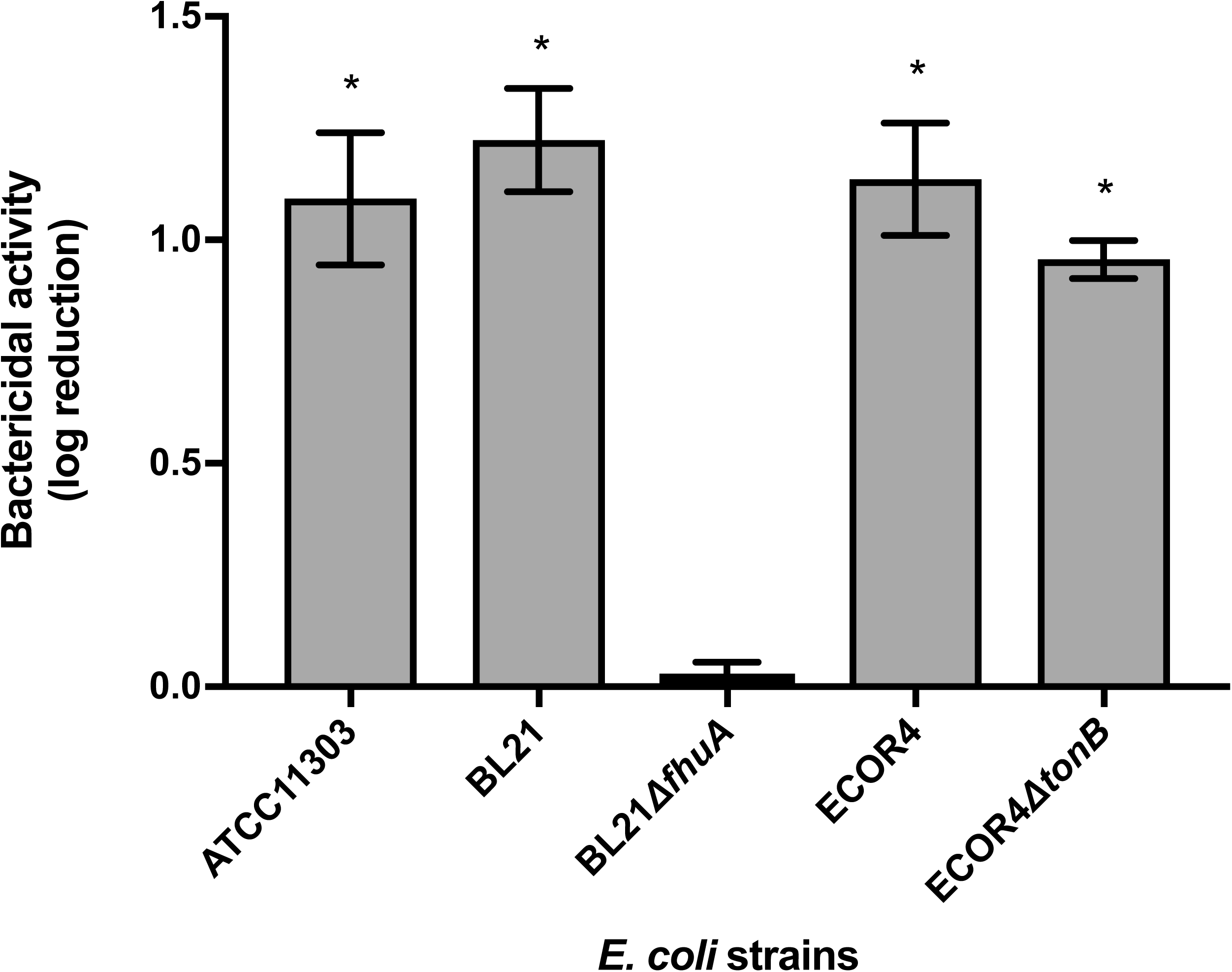
Bactericidal activity of the Innolysin against *E. coli*. Average logarithmic reductions of different *E. coli* strains, after incubation with Innolysin#6 for 30 min at 20°C, compared to the untreated cells. The average reduction was calculated based on triplicates of three independent experiments. * Significant reduction at P < 0.05.

### Morphological changes in cells treated with Innolysin

Transmission Electron Microscopy (TEM) analysis was performed to determine the effects of Innolysin#6 on cell morphology and viability. The morphology of both *E. coli* BL21 and *E. coli* BL21*ΔfhuA* was visualized after incubation with Innolysin#6 for 15 min and compared to untreated cells (Fig. 5). Almost all untreated cells were intact with normal cell envelope morphology. In contrast, treatment of *E. coli* BL21 with Innolysin#6 led to cell integrity damage in the majority of cells with cytosol leakage occurring mainly at the poles. Furthermore, periplasmic space appeared to widen with the outer membrane disconnecting from the inner membrane and cell debris could be detected likely due to cell lysis. This dramatic effect on cell morphology was not observed on *E. coli* BL21*ΔfhuA* treated with Innolysin#6, where only a low increase in damaged cells was noticed compared to the untreated cells. These observations demonstrate that Innolysin#6 acts rapidly by interfering with membrane integrity, while presence of FhuA is required for Innolysin to exert high antimicrobial activity.

**FIG 5:**
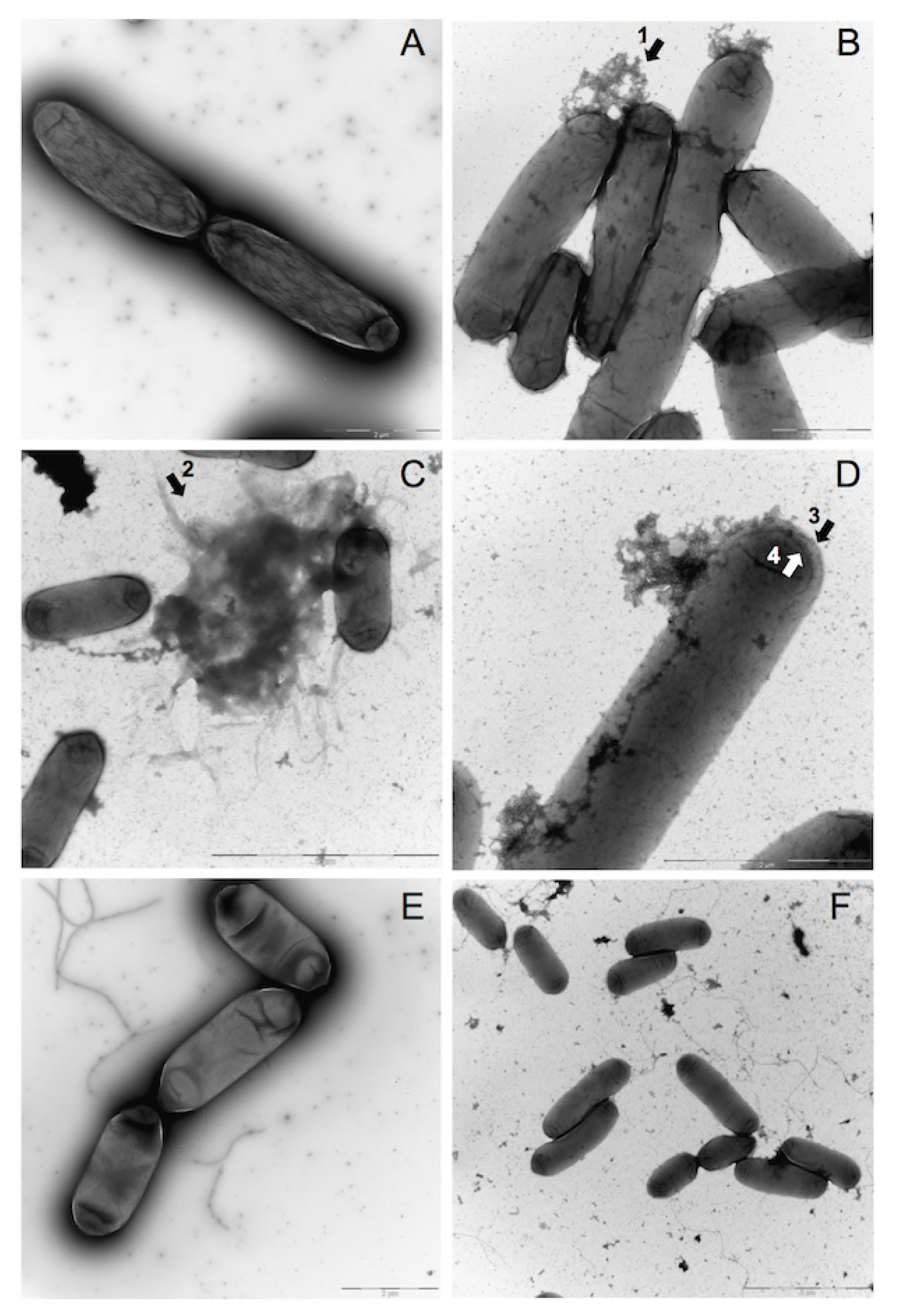
Morphological changes of *E. coli* treated with Innolysin#6. The effect of Innolysin#6 on BL21 (Fig 5B-D) and BL21*ΔfhuA* (F) was compared to the untreated BL21 (Fig 5A) and BL21*ΔfhuA* (Fig 5E), respectively. Cytosol leakage mainly at poles (Fig. 5B, arrow 1); cell debris likely due to cell lysis (Fig. 5C, arrow 2) and separation of the outer membrane (Fig. 5D, arrow 3) and inner membrane (Fig. 5D, arrow 4) were observed in the majority of BL21 cells treated with the Innolysin. Such dramatic changes were not observed in BL21*ΔfhuA* cells after incubation with Innolysin#6 (Fig. 5F).

### Innolysin targets FhuA homologs of other than *E. coli* species

To investigate whether the constructed Innolysin#6 could target FhuA homologs in other species, we tested the killing activity of the purified engineered protein against *Shigella sonnei, Salmonella* Typhimurium LT2c and *Pseudomonas aeruginosa* PAO1 (Fig. 6). These bacteria carry FhuA homologs with identity ranging between 22.6 and 99.6% to the FhuA of *E. coli* BL21 (see Table S2 in the supplemental material). The Innolysin killed *S. sonnei* (99.6% FhuA identity at the protein level) and *P. aeruginosa* PAO1 (less than 39% FhuA identity), leading to average log reductions in cell number of 1.52 ± 0.14 and 1.03 ± 0.25, respectively. In contrast, no significant reduction of *Salmonella* Typhimurium LT2c was noticed, despite the high similarity between the FhuA homolog in this strain and FhuA of *E. coli* BL21 (76.53% FhuA identity). Alignment of FhuA homologs showed that in comparison to FhuA of *S. sonnei* and *E. coli,* FhuA of *Salmonella* Typhimurium LT2c lacks 17 amino acids which are part of the surface-exposed L4 loop of *E. coli* FhuA (Fig. 7) (8). *P. aeruginosa* PAO1 lacks FhuA, however, it harbors other outer membrane proteins involved in the bacterial iron-uptake, including FpvA, FpvB, FptA, FiuA and FoxA (23). Alignment of *E. coli* FhuA with these proteins revealed a low percent identity ranging between 22.61 and 38.52% (see Table S2). Notably, the amino-acid sequence responsible for L4 loop formation of FhuA aligned to parts of the outer membrane proteins FptA and FpvA of *P. aeruginosa* (Fig. S1) which, based on their tertiary structure predictions, form surface-exposed loops (24, 25). This observation suggests that such loops may interact with the Innolysin. In summary, we conclude that the Innolysin can target FhuA homologs in other than *E. coli* species.

**FIG 6:**
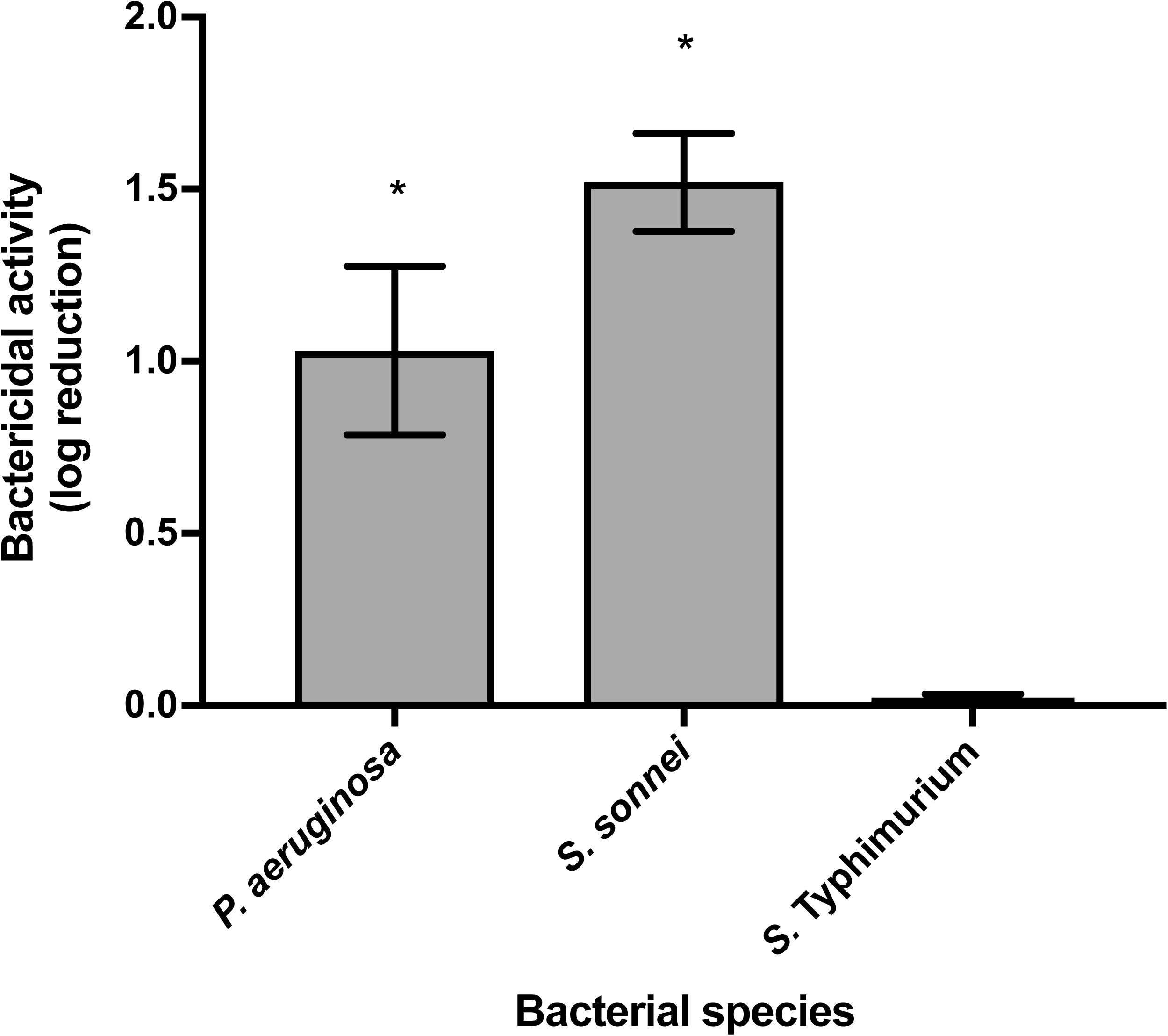
Bactericidal spectrum of the Innolysin. Average logarithmic reductions of different species, after incubation with Innolysin#6 for 30 min at 20°C, compared to the untreated cells. The average reduction was calculated based on triplicates of three independent experiments. * Significant reduction at P < 0.05.

**FIG 7:**
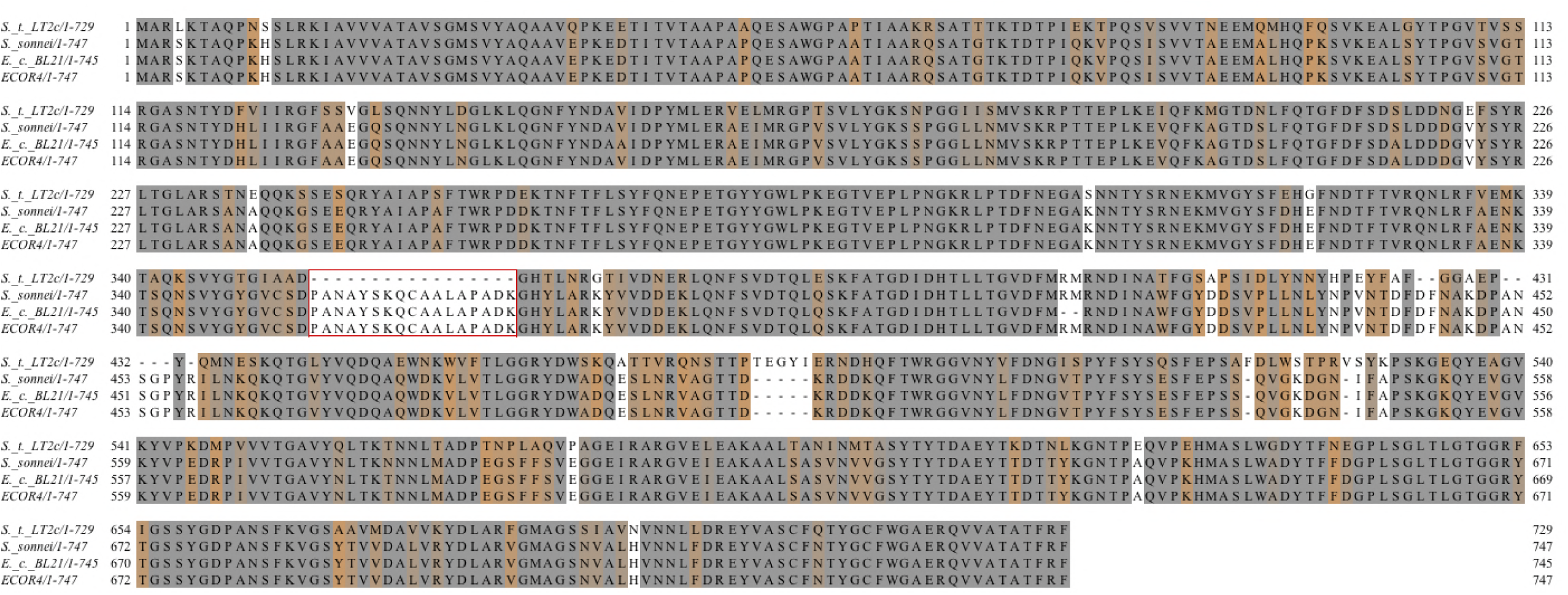
Conservation of FhuA homologs in Gram-negative bacteria. FhuA of *E. coli* BL21 (WP_000124402.1) was used as a reference to search for homologous proteins in other than *E. coli* species. FhuA of *Shigella sonnei* (WP_094317049.1) and *Salmonella* Typhimurium LT2c (NP_459196.1) are aligned with FhuA of *E. coli* BL21 (WP_000124402.1). FhuA protein sequence of *E. coli* ECOR4 (locus tag ESC_AA8779AA_AS_03052, PRJEB2879) is also aligned with FhuA of *E. coli* BL21 to confirm the high conservation between the FhuA of both *E. coli* strains that were used. Amino acids are colored based on the level of conservation with transition of the orange to grey to represent increased conservation. A 5.5 threshold was used for illustrating the conservation of homologs by Jalview sequence alignment tool. The red box highlights the amino acids responsible for the formation of the surface-exposed part of loop L4 of FhuA (WP_000124402.1) in *E. coli* BL21.

## DISCUSSION

Phages have developed unique and complex mechanisms to infect and lyse bacteria. In the first stage of infection, phages utilize receptor binding proteins (RBPs) to target specific bacterial host receptors on the cell surface, whereas at later stages phage endolysins degrade the peptidoglycan layer inducing lysis and progeny release. Here, we exploit the binding capacity of a phage RBP to enable an endolysin to bypass the outer membrane of Gram-negative bacteria and to gain access to peptidoglycan. As proof of concept, we have utilized the phage T5 endolysin and its receptor binding protein Pb5 and constructed twelve Innolysins by fusing the endolysin with the whole Pb5 or with only the binding domain of Pb5 in different orientations, with or without linkers. The majority of the novel Innolysins maintained their muralytic activity and Innolysin#6 also displayed antimicrobial activity against *E. coli.* Our work expands the concept of developing alternative antimicrobials based on enzyme fusions and demonstrates that a phage RBP efficiently can be used to target antimicrobial enzymes selectively to Gram-negative bacterial species.

Here we showed that the bactericidal activity of Innolysin#6 consisting of T5 endolysin, a small linker and the binding domain of Pb5, is dependent on FhuA receptor of phage T5 (5, 6). Similar to phage T5 binding and DNA injection (22, 26, 27), the antimicrobial activity of Innolysin#6 is independent of energy provided by TonB, indicating that Innolysin#6 is passively transported to the peptidoglycan. Microscopic analysis revealed that Innolysin#6 causes cytosol leakage mainly from the bacterial poles of wild type *E. coli* encoding FhuA. Interestingly, several phages have been shown to preferentially adsorb at the bacterial poles, including *E. coli* phage φ80 that uses FhuA as receptor (28). Thus, Innolysin#6 may bind to FhuA of *E. coli* in a similar manner as phage T5 causing specific lysis of cells carrying the FhuA receptor of Pb5.

Since FhuA is the binding target of Innolysins and works as a transport channel for uptake of ferrichrome, it is tempting to speculate that Innolysin is transported passively through the channel. Yet, it has been shown that Pb5 binding to FhuA does not allow the channel to open (5, 9) and similarly binding of Innolysin may not provide the conformational changes needed for the FhuA channel to open. In addition, the size of the unplugged channel (2.5 nm) appears to be too narrow for the Innolysin (67.62 kDa) to pass (29), indicating that partial unfolding is required for the Innolysin to overcome the size restraint by FhuA. Lukacik and co-workers showed that a hybrid lysin consisted of the phage T4 lysozyme and the binding domain of pesticin targeting FyuA could reach the peptidoglycan (15). But similar to Innolysins, the dimensions of the FyuA pore do not permit passage of the hybrid T4 lysozyme without unfolding. Yet, it was demonstrated that unfolding of the enzymatic domain is not required for activity (15), indicating that such hybrid endolysins may overcome the outer membrane by other means than passing through the targeted channels.

Alternatively, Innolysins may access the peptidoglycan by interfering with the membrane integrity. FhuA has been proposed to work as an anchor for phage T5 by an irreversible binding of Pb5 to the FhuA receptor (10). In Innolysins we combine Pb5 with phage T5 endolysin that is a globular endolysin carrying a single enzymatically active domain with limited intrinsic antimicrobial activity (30). Thus, we equipped a globular endolysin with a binding domain using a phage RBP and transformed a globular endolysin into a novel engineered modular endolysin. In comparison to globular endolysins, modular endolysins composed of catalytic domain(s) and a cell wall binding domain display up to five times higher enzymatic activity (31, 32). Thus, by irreversible binding to FhuA (5), Pb5 may bring the Innolysin in close proximity to the outer membrane, enhancing the antibacterial activity of the endolysin. The binding of the Innolysin could interfere with phosphates in the outer membrane lipopolysaccharides and displace the stabilizing Mg^2+^/Ca^2+^ ions due to the high isoelectric point of phage T5 endolysin (7.91), thus destabilizing the ionic forces in the outer membrane (33). A similar mode of action has been described for Artilysins, endolysins engineered with polycationic or amphipathic peptides that destabilize the outer membrane and target the endolysins to peptidoglycan (14). Furthermore, we found that a linker between the T5 endolysin and Pb5_1-488_ components is essential for bactericidal activity. Similarly, modular endolysins contain flexible interdomain linkers, which are important for enzyme activity (34). Thus, the linker may be necessary for the Innolysin to properly bind to the receptor FhuA, interfere with the membrane integrity and overcome the outer membrane.

FhuA is conserved in bacterial species including *Escherichia, Shigella* and *Salmonella* and is structurally homologous to other Ton-B dependent outer membrane receptors involved in bacterial iron uptake (35). The Innolysin showed antimicrobial activity towards *Shigella sonnei* and *Pseudomonas aeruginosa*, whereas *Salmonella* Typhimurium LT2c was not sensitive to the Innolysin. While *S. sonnei* and *P. aeruginosa* encode homologs or equivalents of FhuA and share a conserved L4 loop, shown to be the binding target of phage T5 (22, 26), *Salmonella* FhuA lacks the L4 loop. Thus, the L4 loop of FhuA may be required for the antimicrobial activity of the Innolysin, enabling efficient binding of Pb5 to its cognate receptor. However, since FhuA is a phage receptor, common phage adsorption blocking mechanisms, such as receptor masking by lipopolysaccharides or downregulation of receptor expression (36, 37) could also play a role in Innolysin resistance in *Salmonella*.

In this study, we used the binding specificity of the phage T5 receptor binding protein to its cognate FhuA receptor to deliver an endolysin across the outer membrane and target the peptidoglycan. Compared to bacteriocin-derived fusions with endolysins, the natural diversity of phage RBPs greatly expands the potential of designing novel antimicrobials targeting a wide range of Gram-negative bacterial species. As opposed to some bacteriocins and traditional antibiotics, Innolysins can target non-actively growing bacteria, since they bypass the outer membrane passively (38). Additionally, Innolysins can be designed to target important virulent determinants such as OmpX in *Escherichia coli* and Ail in *Yersinia pesti*s, in order to modulate virulence (39, 40). Although further experiments are needed for optimizing the activity of such antimicrobials and for elucidating the exact mechanism by which Innolysins bypass the outer membrane, our findings provide a promising approach to design novel therapeutic agents with a highly customizable bactericidal spectrum.

## MATERIAL AND METHODS

### Bacterial strains

*E. coli* BL21-CodonPlus-(DE3)-RIL (Agilent Technologies) was used for expression of recombinant proteins. Inhibition assays were conducted with *E. coli* ATCC11303 (Leibniz Institute). *E. coli* BL21 (DE3) (Agilent Technologies), *E. coli* ECOR4 (STEC Center at Michigan State University), *Shigella sonnei* (41), *Pseudomonas aeruginosa* PAO1 (42) and *Salmonella Typhimurium* LT2c (43) were used for the bactericidal assay. Deletions of the *fhuA* and *tonB* genes on the chromosomes of *E. coli* BL21 (DE3) and *E. coli* ECOR4, respectively, were previously generated by the Lambda red recombination system as described before (44). Briefly, the kanamycin cassette from plasmid pKD4 was amplified (*Pfu* polymerase; Thermo Fisher Scientific, Waltham, MA) with oligonucleotides carrying ∼39 nucleotide extensions homologous to regions adjacent to the target gene (custom primers, Table S1). The amplicon was purified with QIAquick PCR Purification Kit (Qiagen) and electroporated into competent cells. Kanamycin-resistant clones were selected and insertion of the kanamycin cassette was verified by PCR.

### Cloning of constructs

Genomic material was purified from phage T5 (Leibniz Institute) and used as a template for amplification (*Pfu* polymerase; Thermo Fisher Scientific, Waltham, MA) of the genes encoding T5 endolysin (YP_006868.1) or Pb5 (YP_006985.1) with specific primers (Table S1). The linkers were created as primer cassettes. The specific primers were mixed, heated to 95°C and gradually cooled down for hybridization. Each assembly encoding the different fragments of the fusion proteins was constructed by Type IIs cloning. Amplicons or primer cassettes (each 50 ng/µl) were mixed with 100 ng/µl pNICBsa4, in which the N-terminal His-tag was exchanged for a C-terminal His-tag, and BsaI and T4 DNA ligase were added. The mixture was incubated alternatingly at 37°C and 16°C (50 cycli) and heat inactivated at 80°C. Transformation of *E. coli* BL21-CodonPlus-(DE3)-RIL was conducted and transformants were selected in the presence of kanamycin (100 μg/ml) and chloramphenicol (50 μg/ml).

### Protein expression and purification

Recombinant proteins were simultaneously expressed in a 96 deep-well plate. Freshly transformed colonies were resuspended in 500 μl of auto-induction medium (93% ZY medium, 0.05% 2M MgSO_4_, 2% 50x 5052, 5% 20x NPS) and incubated at 37 °C for 5 hours, followed by 16 °C for 40 hours, both at 900 rpm. Cells were spun down by centrifugation (3,200 × *g*, 30 min, 4°C) and the supernatants were removed. Cell pellets were lysed by exposure to chloroform vapor, by placing the deep-well plate upside-down in a glass petri dish containing chloroform-saturated filters for 2 h. Cell lysates were resuspended in 20 mM HEPES-NaOH (pH 7.4), and 1 U DNase I, followed by incubation (100 × rpm, 1 hour, 30 °C). Insoluble fractions of cell lysates were removed by centrifugation (3,200 × *g* for 30 min, at 4°C) and the supernatants containing the soluble fractions of the lysates were screened for both muralytic and inhibitory activities.

For large-scale expression of the Innolysin#6, a 1 L expression culture (LB medium) was induced with 1 mM isopropyl-beta-D-thiogalactopyranoside (Thermo Fisher Scientific) in the mid-logarithmic phase (OD_600_=0.6). Incubation of the culture was followed at 16°C for eighteen hours at 120 rpm. Cells were harvested by centrifugation (8,000 × *g*, 10 min, 4°C) and the cell pellet was resuspended in 10 ml of lysis buffer (20 mM NaH_2_PO_4_-NaOH, 0.5 M NaCl, 50 mM imidazole, pH 7.4), followed by sonication (Bandelin Sonopul HD 2070 homogeniser) with 10 bursts of 30 sec (amplitude of 50%) with 30 sec intervals. Protein lysate was double-filtered using filters with pore size of 0.22 μm. His GraviTrap™ gravity flow columns (GE Healthcare) were used for His-tagged protein purification according to the manufacturer’s instructions. Buffer exchange was performed with 20 mM HEPES-NaOH (pH 7.4) by using Amicon^R^ Ultra - 4 centrifugal filters with 50 kDa cutoff (Merck Millipore) and protein concentration was measured by Qubit™ Protein Assay Kit (Q33211) in Qubit 2.0 Fluorometer (Invitrogen, Q32866).

### Muralytic assay

Analysis of the muralytic activity was conducted as described before (14, 45) using outer membrane permeabilized *P. aeruginosa* PAO1 cells as substrate. Briefly, exponentially growing cells (OD_600_ = 0.6) were harvested by centrifugation (3,200 × *g*, 30 min, 4°C) and permeabilized by resuspension in chloroform-saturated 0.05 M Tris-HCl (pH 7.7) and gentle shaking for 45 min. To remove chloroform traces, cells were washed twice with phosphate-buffered saline (PBS, pH 7.4) and further concentrated to OD_600_ = 1.5. 30 μl of the soluble lysate fractions was added on top of 270 μl of the substrate. Supernatants of lysates of cells expressing the phage T5 endolysin or carrying an empty vector (pVTSD2) were used as positive and negative controls, respectively. Turbidities were measured spectrophotometrically at 655 nm every 30s for one hour by a Microplate Reader 680 system (Bio-Rad). Muralytic activities were calculated by a previously described standardized method (46).

### Growth inhibition assay

*E. coli* ATCC11303 cells were used for the growth inhibition assay. Overnight cell cultures prepared in Mueller Hinton (MH) broth were adjusted to OD_600_= 0.1 in 2xMH and further 100-fold diluted in 2xMH. 50 µl of the cell suspension was mixed with 50 μl of the soluble lysate fraction. Art-175 (0.1 mg/ml) and the soluble lysate fraction of cells carrying an empty vector were mixed with cells as positive and negative controls, respectively. Endpoint measurement was performed spectrophotometrically at 655 nm after exactly 18 h incubation at 37°C. All experiments were done in triplicate.

### Antibacterial assay

Gram-negative, exponentially growing cells (OD_600_=0.6) were diluted 100-fold in 20 mM HEPES-NaOH (pH 7.4). 100 μl of the diluted cell suspensions were mixed with 100 μl of the pure Innolysin at a final concentration of 0.2 mg/ml. 100 μl of 20 mM HEPES-NaOH (pH 7.4) was added to the cells as a negative control. The samples were incubated for 30 min at 20°C, and appropriate cell dilutions were plated on LB (Lysogeny Broth) agar plates in triplicate. After overnight incubation at 37°C, colony forming units (CFU) were counted and cell concentrations (CFU/ml) were calculated. Experiments were conducted three independent times. The antibacterial activity was determined based on the difference of the average logarithmic cell concentrations of the treated samples compared to the negative control.

### Transmission Electron Microscopy

1 ml of exponentially growing cells were harvested by centrifugation (10000 rpm, 5 min) and resuspended in 200 μl of HEPES buffer (pH 7.4). This washing procedure was repeated 3 times, after which the cell pellets were resuspended in either 50 μl of HEPES buffer or the Innolysin#6 and incubated for 15 minutes at 20°C. Samples were negatively stained with 2% uranyl-acetate on glow-discharged continuous carbon-coated 300-mesh copper grids (EM Resolutions Ltd). Transmission electron microscopy was performed on a Philips CM100 (Tungsten emitter) electron microscope operating at 100 kV. Images were recorded on a side-mounted Olympus Veleta (2048 x 2048 pixels) charge-coupled device camera via the iTEM software.

### Bioinformatic analysis

To identify FhuA homologs, we used the FhuA protein sequence of *E. coli* BL21 (WP_000124402.1) to search homologous proteins in the National Center for Biotechnology Information (NCBI) genome database through BLASTP (47). Homologous proteins with different levels of identity in other than *E. coli* species were selected and aligned by the multiple sequence alignment Clustal Omega (48), through which the percent identity was obtained. To further identify the conserved homologs of the proteins we used Jalview sequence alignment tool (49).

### Statistical analysis

Analysis of the data was conducted by using GraphPad Prism 7 software (Version 7.0d). For muralytic assays, the activity of each soluble lysate fraction was tested in triplicate and means of activity was compared with the average activity of soluble lysate fraction of cells carrying an empty vector. The significance of muralytic activity was assessed with Unpaired-Samples t-test using 95% confidence interval for the mean difference. The same software was used to analyze results from the growth inhibition assays, using the optical density (OD_655_) of the bacterial growth after incubation with the soluble lysate fraction of each protein compared to the optical density of cells grown after treatment with soluble lysate fraction of cells carrying an empty vector. For the bactericidal assay, all bacterial counts were converted to log-scale and means and standard deviations were calculated afterwards. Bactericidal activity was tested in triplicate in three independent experiments. Decimal reductions of cells were calculated by the difference between the average logarithmic concentrations of cells treated with Innolysin#6 and untreated cells, after incubation. The significance of the decimal reductions of cells was assessed with Paired-Samples t-test using 95% confidence interval percentage.

## ACKNOWLEDJMENTS

This work was supported by the Danish Council for Independent Research (4184-00109B). We sincerely thank Lysando AG for supplying Artilysin-175 and our colleague Stephen Ahern for supplying the *fhuA* and *tonB* mutants of *E. coli*. Furthermore, we would like to thank Yilmaz Emre Gencay and Hans Gerstmans for fruitful discussions.

## SUPPLEMENTAL MATERIAL

**FIG S1 Alignment of *E. coli* BL21 FhuA with homologous proteins of *P. aeruginosa*.**

FiuA (NP_249161.1), FoxA (NP_251156.1), FptA (NP_252911.1), FpvA (NP_251088.1) and FpvB (NP_252857.1) of *Pseudomonas aeruginosa* PAO1 were aligned with *E. coli* BL21 FhuA (WP_000124402.1). Amino acids are colored based on the level of conservation with transition of orange to grey to represent higher conservation. Conservation with no threshold was used by Jalview sequence alignment tool. The red box highlights the alignment of *P. aeruginosa* homologous proteins to the amino acid sequence of FhuA responsible for the formation of the surface-exposed part of loop L4 in *E.coli* BL21.

**Table S1: Primers used for PCR amplification in cloning**

For generating mutations in *E. coli* strains, primers were designed with regions of homology to the target genes (indicated in bold). To engineer recombinant proteins, *oad* and *lys* genes of phage T5 encoding Pb5 and phage T5 endolysin, respectively, were used for fusion, whereas linkers L1: AGAGAG and L2: GAGAGAGAGAGAGA were created as primer cassettes. The recognition sites of BsaI (highlighted in grey) were used for cloning into pNICBsa4 vector.

**Table S2: Percent matrix identity of BL21 FhuA homologs**

Percent identity between homologous proteins and *E. coli* BL21 FhuA (WP_000124402.1) was calculated by sequence alignment Clustal Omega. Full-length FhuA protein sequences of *E. coli* BL21 (WP_000124402.1), *E. coli* ECOR4 (locus tag ESC_AA8779AA_AS_03052, PRJEB2879), *Shigella sonnei* (WP_094317049.1) and *Salmonella* Typhimurium LT2c (NP_459196.1) were used for constructing the identity matrix. Furthermore, homologous proteins of *Pseudomonas aeruginosa* PAO1, including FiuA (NP_249161.1), FoxA (NP_251156.1), FptA (NP_252911.1), FpvA (NP_251088.1) and FpvB (NP_252857.1), were tested for the identity level compared to *E. coli* BL21 FhuA (WP_000124402.1).

